# A molecular switch for stress-induced activation of retrograde mitochondrial transport

**DOI:** 10.1101/2024.09.13.612963

**Authors:** Christina Gladkova, Maria G. Paez-Segala, William P. Grant, Samuel A. Myers, Yuxiao Wang, Ronald D. Vale

## Abstract

The cellular distribution of mitochondria in response to stress and local energy needs is governed by the relative activities of kinesin and dynein. The mechanism for switching between these two opposite polarity microtubule motors remains unknown. Here, we coupled a novel cellular synthetic cargo transport assay with AlphaFold2-guided mutagenesis to identify a regulatory helix in the mitochondrial adaptor protein (TRAK) that mediates switching between kinesin- and dynein-driven transport. Differences in the helix sequence explain why two near-identical TRAK isoforms transport mitochondria in predominantly opposite directions. Phosphorylation of the regulatory helix by stress-activated kinases causes the activation of dynein and dissociation of kinesin. Our results reveal a molecular mechanism for coordinating the directional transport of mitochondria in response to intracellular signals.

## INTRODUCTION

Membrane-bound organelles are transported by kinesins and cytoplasmic dynein along microtubules, which, in most cells, are organized as a polarized network with their minus-ends anchored at the centrosome near the cell center and plus ends oriented towards the cell periphery. Most organelles exhibit bidirectional transport, with episodes of kinesin-driven (plus-end-directed; anterograde) and dynein-driven (minus-end-directed; retrograde) movement (*1*). The distribution of a particular organelle is governed by the net activity of the two motors. Organelles such as mitochondria tend to move in the anterograde or retrograde direction at speeds characteristic of the maximal velocity of kinesin or dynein motors (*2–4*), suggesting that kinesin and dynein on an organelle are not continuously opposing one another, and that regulatory mechanisms for coordinating motor binding and/or activity likely exist.

Mitochondria constitute perhaps the best studied example of bidirectional organelle transport (*5, 6*). Mitochondrial transport by kinesin-1 supports energetically demanding peripheral processes such as cell outgrowth or synaptic activity (*7, 8*), while retrograde transport by dynein towards the centrosome is necessary for turnover of damaged mitochondria (*9*). TRAK, a multi-functional motor adaptor protein (*10*), recruits kinesin-1 and dynein (*11–13*), and also binds to mitochondria via an outer membrane receptor Miro (*13–15*) (Fig. 1A). TRAK has several features that enable it to interact with the active dynein complex and that are conserved in related adaptor proteins (*16*): a CC1-box that interacts with the dynein light intermediate chain (dynein LIC) (*17, 18*), a Spindly motif that binds the dynein activator dynactin (*19*), and a long coiled-coil domain that is sandwiched between the dynein and dynactin complexes in their active conformation (*16–18*). The TRAK coiled-coil also contains the binding site for kinesin-1, although the precise way in which TRAK binds kinesin-1 is not known (*20–22*). Recent in vitro studies have examined how motor transport is coordinated by TRAK, but emerged with two different models, suggesting either co-dependence (*23*) (one motor aiding the other), or mutual exclusivity (*24*) of kinesin and dynein activity.

**Fig. 1.**
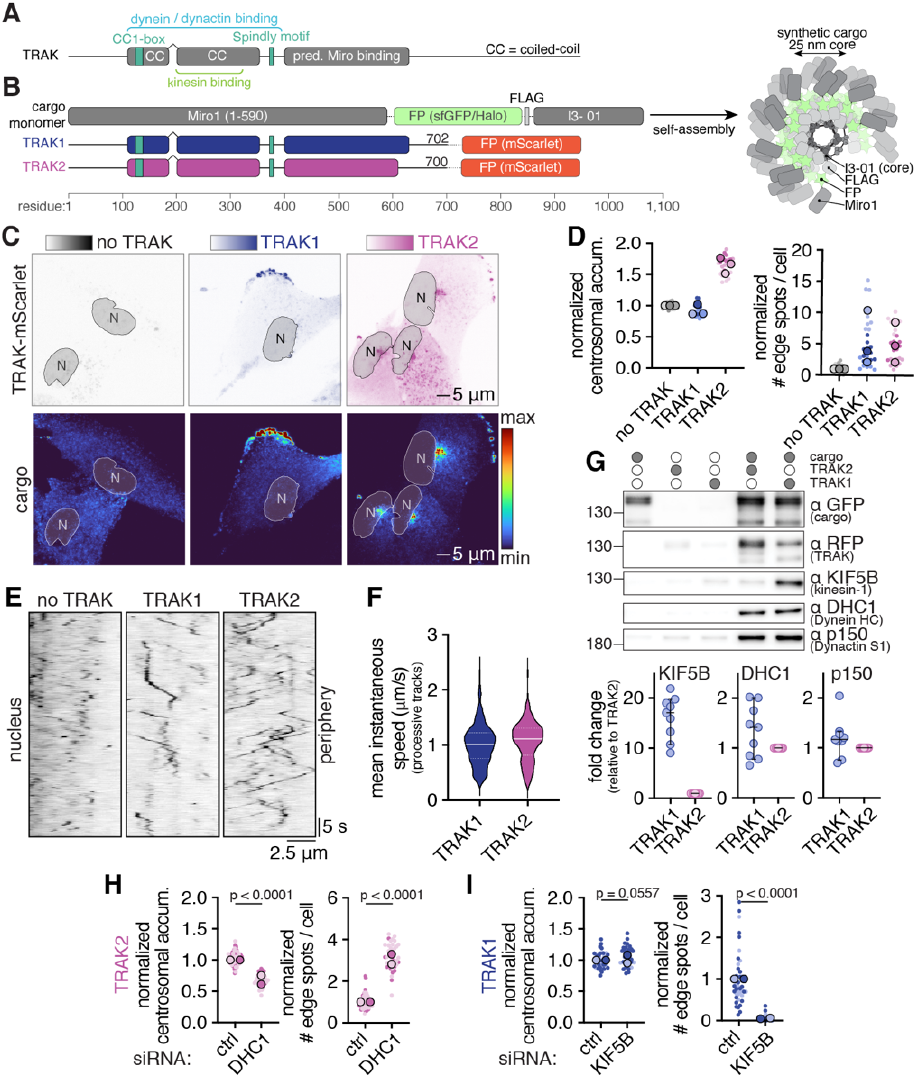
Microtubule-based transport of synthetic mitochondrial cargo. (**A**) Common domain organization for TRAK1 and TRAK2 with interaction sites annotated based on prior work. (**B**) Generating synthetic mitochondrial cargo by lentiviral expression of Miro1/FP/FLAG/I3-01 (FP = Superfolder GFP (sfGFP), or a Halo-Janelia Fluor Dye conjugate) and TRAK/mScarlet fusion proteins in hTERT RPE-1 cells. 60 copies of the I3-01 fusion assemble to form the synthetic cargo (*32*). (**C**) Representative maximum intensity projection images of cells expressing TRAK (top) and synthetic cargo (bottom). (**D**) Phenotype quantification of synthetic cargo accumulation from cells as in (C) imaged at lower resolution. Data points correspond to normalized median values calculated for a field of view with ∼100 cells, color-matched means from repeated experiments are superimposed. See fig. S3 and Methods for analysis description. (**E**) Kymographs showing processive motility of PA-JF646 labelled synthetic cargo after photoactivation. Kymographs were generated by reslicing time series shown in Movie 2 as indicated in fig. S2C. (**F**) Mean instantaneous speed averaged over the length of processive tracks (maximum distance travelled >3.0 μm). This speed was 1.02 ± 0.02 μm/s for TRAK1 (n = 21 cells, 573 tracks), and 1.07 ± 0.02 μm/s for TRAK2 (n = 47 cells, 270 tracks). See fig. S2C and Methods for analysis description. (**G**) Top: Immunoblot analysis of the motor-adaptor components associated with FLAG-purified cargo from indicated cell lines. Bottom: quantification of relative motor protein levels normalized to TRAK2. Error bars show the median and 95% confidence interval; n = 9 for all conditions. (**H**) Quantification of cargo accumulation in cells expressing TRAK2 following DHC1 RNAi from two biological replicates. Representative maximum intensity projection images are shown in fig. S2G. For perinuclear accumulation n(ctrl)=74; n(DHC1)=39; p<0.0001 (two-tailed t-test, t=17.75, df=111). For peripheral accumulation n(ctrl)=72; n(DHC1)=38; p<0.0001 (two-tailed t-test, t=2.936, df=47). (**I**) Quantification of cargo accumulation in cells expressing TRAK1 following KIF5B RNAi from two biological replicates. Representative maximum intensity projection images are shown in fig. S2H. For perinuclear accumulation n(ctrl)=44; n(DHC1)=33; p=0.0557 (two-tailed t-test, t=1.943, df=75). For peripheral accumulation n(ctrl)=41; n(KIF5B)=31; p<0.0001 (two-tailed t-test, t=8.105, df=70).

Vertebrates have two TRAK adaptor genes, TRAK1 and TRAK2, that are ∼75% similar in the motor binding region (fig. S1). Both TRAKs bind dynein/dynactin equally well; however, TRAK1’s affinity for kinesin is much greater compared to TRAK2 (*11, 24*). In hippocampal neurons, TRAK1 preferentially transports cargos into the axon, while TRAK2 predominately drives transport into dendrites (*11, 25*). Previous cellular studies also found that TRAK-mediated transport is modulated in response to different cues such as cell cycle status (*26*), glucose availability (*27, 28*), and the presence of reactive oxygen species (*29*). However, the differences between the two TRAKs, how TRAK switches between kinesin- and dynein-driven transport, and how motor switching can be influenced by intracellular signals remain poorly understood.

## RESULTS

### A synthetic cargo recapitulates mitochondrial transport

The complex interactions of mitochondria with actin, microtubules, and other organelles as well as their fission and fusion dynamics pose challenges for isolating and studying the coordination of microtubule-based motors in cells (*5, 30, 31*). Therefore, we sought to design a simplified cell-based assay that reports specifically on regulatory mechanisms governing microtubule-based transport of mitochondria. To this end, we constructed a “synthetic cargo” based on an engineered protein (I3-01) that self-assembles into ∼25 nm dodecahedral nanocages (*32*). To study mitochondrial transport in hTERT RPE-1 cells, we expressed I3-01 monomers fused to the cytoplasmic domain of Miro1 (the TRAK mitochondrial adaptor), a fluorescent protein for visualization, and a FLAG tag for biochemical characterization and co-expressed TRAK1 or TRAK2 constructs fused to the fluorescent protein mScarlet (Fig. 1B).

When expressed in hTERT RPE-1 cells, sfGFP-labelled synthetic cargos were uniformly distributed throughout the cytoplasm, suggesting that endogenous TRAK is insufficient for their robust transport. However, co-expression of TRAK1 or TRAK2 led to a dramatic, microtubule-dependent synthetic cargo relocalization (Fig. 1C; fig. S2A, Movie S1). Expression of TRAK1 resulted in the peripheral accumulation of synthetic cargo, consistent with kinesin-driven transport. Expression of TRAK2 resulted in predominant accumulation of synthetic cargo at the perinuclear region, consistent with dynein-driven transport. We developed a pipeline to quantitate this steady-state perinuclear and peripheral accumulation of cargo in hundreds of cells (Fig. 1D, fig. S3). To investigate the dynamics of cargo accumulation, we swapped the sfGFP with a HaloTag conjugated with the photoactivatable JF-646 Halo dye (*33*). We observed unidirectional transport of individual particles consistent with their bulk localizations (Fig. 1E, fig. S2 B-D, Movie S2), at rates similar to those reported for kinesin- and dynein-driven transport of mitochondria in vivo (*2*) (Fig. 1F, fig. S2E).

The distinct preferences for synthetic cargo transport by TRAK1 (kinesin-driven) and TRAK2 (dynein-driven) mirror similar findings for mitochondria (*34*) (fig. S2F). Consistently with previous studies (*11, 24*), dynein and dynactin associated equally well with TRAK1- and TRAK2-driven synthetic cargo, while kinesin was ∼20 fold enriched in TRAK1-driven synthetic cargo (Fig. 1G). Next, we investigated whether depletion of the dominant motor by RNAi would allow the opposite polarity motor to take over. In the case of TRAK2, dynein knockdown shifted the predominant cargo accumulation from the perinuclear region to the cell periphery (Fig. 1H, fig. S2G), indicating that kinesin can take over transport when dynein is depleted. In the case of TRAK1, kinesin knockdown predominantly led to diffuse cargo localization without centrosomal accumulation (Fig. 1I, fig. S2H), suggesting that the dynein/dynactin bound to TRAK1 was inactive, even in the absence of competing kinesin.

In summary, our synthetic cargo assay recapitulates known features of mitochondrial transport, and suggests that TRAK2 has the ability to switch between kinesin- and dynein-mediated transport, while TRAK1 exists in a repressed state that can associate with, but not activate dynein/dynactin.

### AlphaFold insight into TRAK-motor interactions

We next employed AlphaFold2-multimer to better understand the mechanisms of TRAK association with kinesin and dynein and repression of dynein/dynactin by TRAK1 (*35*). We generated a prediction for the N-terminal coiled-coil dimer of TRAK1, as well as model for its interaction with kinesin (Fig. 2A). The predictions for the analogous TRAK2 complexes were near-identical (fig. S4A), and equally confident (fig. S5).

**Fig. 2.**
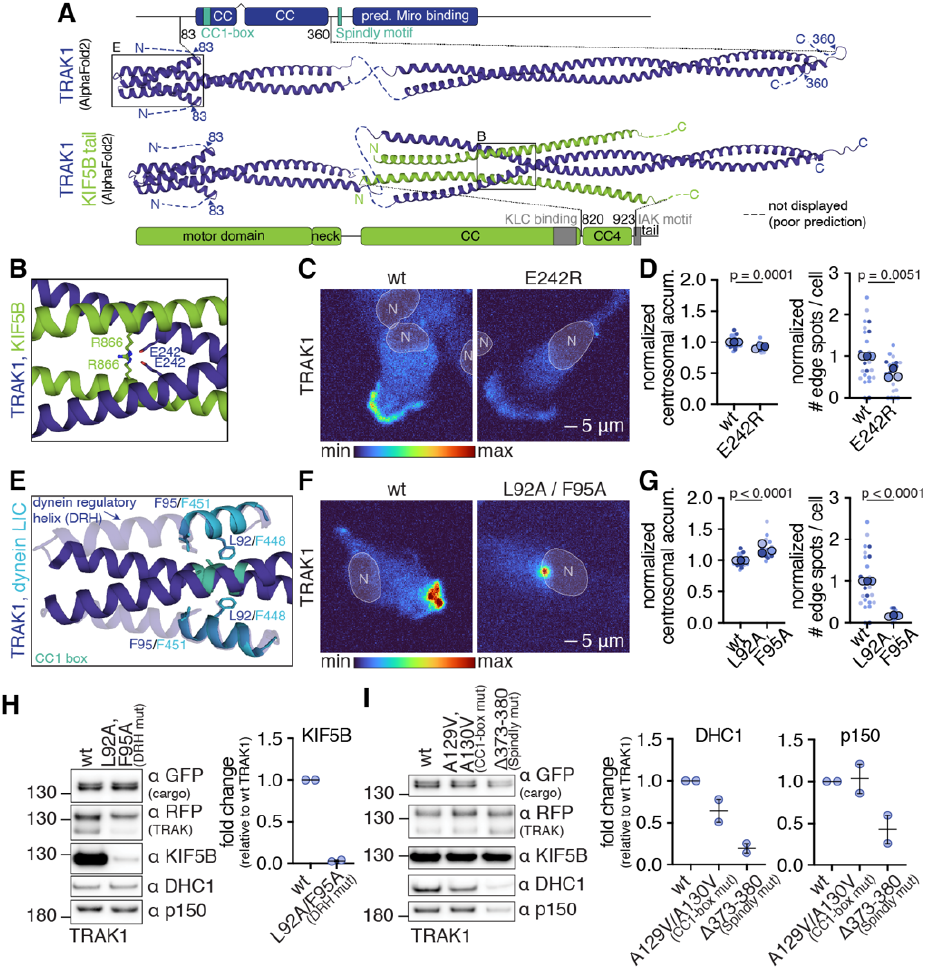
An AlphaFold2-derived model for TRAK-motor interactions. (**A**) AlphaFold2 predictions for TRAK1 (blue). Top: N-terminal coiled-coil dimer (residues shown 83-360). Bottom: N-terminal coiled-coil dimer in complex with the CC4 region of kinesin-1 (green, residues shown 820-923). For clarity, the TRAK1 coiled-coil is split into two separately aligned sections at a site of disorder (see fig. S5). (**B**) Inset from the kinesin complex prediction in (A) showing residues E242 in TRAK1 and R866 in kinesin-1 that are predicted to form a buried salt-bridge. (**C**) Representative maximum intensity projection images of cargo from cell lines expressing wt or E242R TRAK1. (**D**) Phenotype quantification from three biological replicates as shown in (C). For perinuclear accumulation n(wt)=27; n(E242R)=25; p=0.0001 (two-tailed t-test, t=4.107, df=50). For peripheral accumulation n(wt)=26; n(E242R)=23; p=0.0051 (two-tailed t-test, t=2.936, df=47). (**E**) Inset from the TRAK1 coiled-coil prediction in (A) showing the CC1-box of TRAK1 (cyan) with dynein LIC (light blue) superimposed based on a crystal structure of the dynein LIC:BICD2 adaptor complex(*17*) (PDB ID: 6pse). Hydrophobic residues of the N-terminal TRAK1 folded back helix (L92, F95) mimic interfacing dynein LIC residues (F448, F451). (**F**) Representative maximum intensity projection images of cargo from cell lines expressing wt or L92A/F95A TRAK1. (**G**) Phenotype quantification from three biological replicates as shown in (F). For perinuclear accumulation n(wt)=27; n(L92A/F95A)=23; p<0.0001 (two-tailed t-test, t=5.754, df=48). For peripheral accumulation n(wt)=26; n(L92A/F95A)=22; <0.0001 (two-tailed t-test, t=6.039, df=46). (**H, I**) Left: Immunoblot analysis of the motor-adaptor components associated with FLAG-purified cargo from indicated cell lines. Right: quantification of relative motor protein levels normalized to wt TRAK1. Error bars show the median and 95% confidence interval; n = 2 for all conditions.

The AlphaFold2 prediction for the location of the kinesin binding site on TRAK was consistent with previous mapping experiments (*11, 20*), and suggested that the kinesin C-terminal coiled-coil (CC4, aa 820-923) forms a four-helical bundle with the C-terminal portion of the TRAK coiled-coil (aa 205-260). To test this model, we mutated E242, a highly conserved residue that is predicted to form a buried salt-bridge with kinesin (R866), but is solvent exposed in the kinesin-free state (Fig. 2B, fig. S4 B and C). In support of this model, the E242R mutation led to a decrease in kinesin association with both TRAKs (fig. S4 D and E), although the effect was more pronounced for TRAK2, the inherently weaker kinesin binder. For TRAK2, the mutation led to enhanced perinuclear accumulation, consistent with a reduction in kinesin binding (fig. S4F). For TRAK1 the mutation phenocopied the kinesin knockdown result (Fig. 1I), and led to decreased peripheral localization without corresponding gain of perinuclear accumulation (Fig. 2 C and D).

AlphaFold2 also provided insight into a mechanism for dynein/dynactin repression. Specifically, we noted an N-terminal TRAK helix (aa 85-106) that was folded-back to interact with the coiled-coil region of TRAK to form a four-helix bundle in all of our AlphaFold2 predictions for both TRAK1 and TRAK2 (fig. S6A). This interaction mimics the manner in which the dynein LIC interacts with the CC1-box region of the BICD2 dynein adaptor in a crystal structure (*17*) (Fig. 2E), an interaction known to be important for dynein/dynactin motility (*17, 18*). Thus, with the helix in a ‘folded back’ (closed) position, the binding site for dynein LIC should be occluded. To test the AlphaFold2 prediction and the role of the ‘fold-back’ TRAK helix, we generated alanine substitutions for two key hydrophobic residues (F95, L92 in TRAK1 or M95, F92 TRAK2) that mimic the dynein LIC residues L451 and F448 that interact with the CC1-box. Strikingly, the F95A, L92A TRAK1 mutant promoted transport of cargo to the centrosome, the opposite direction to wild-type TRAK1 (Fig. 2 F and G). This result suggests that disrupting the fold-back helix interaction with the CC1-box was sufficient to activate dynein/dynactin transport by TRAK1. Comparable alanine mutations also enhanced perinuclear accumulation in TRAK2 (fig. S6B).

Together, our AlphaFold-guided mutagenesis results provide a model for kinesin binding, and imply that dynein activation requires a structural rearrangement of the folded back helix from a “closed” to an “open” conformation that permits dynein LIC binding to the CC1-box. Because of its role in regulating dynein activity, we use the term “dynein regulatory helix (DRH)” to describe this helix.

### DRH couples dynein activation with kinesin dissociation

Our cellular motility and biochemical pull-down results taken together show that TRAK1, with a primarily “closed” DRH, associates with active kinesin and inactive dynein/ dynactin. TRAK2, with a more “open” DRH, associates with active dynein/dynactin but not kinesin. These findings are consistent with previous in vitro motility studies showing that anterograde-moving TRAK complexes contained both kinesin and inactive dynein/dynactin, whereas retrograde-moving complexes contained active dynein/dynactin only (*24*). To directly test whether dynein activation mediated by DRH opening leads to kinesin dissociation, we activated TRAK1-associated dynein/dynactin using the DRH opening mutant (F95A, L92A), and measured kinesin association. Strikingly, the DRH mutants dramatically decreased kinesin binding to TRAK1 (Fig. 2H), suggesting that dynein LIC binding to the CC1-box generates a conformation of the TRAK/dynein/dynactin complex that is incompatible with binding kinesin. The comparable TRAK2 alanine mutations also decreased kinesin association (fig. S6C).

Since the CC1-box interaction with dynein LIC appears to be predominantly regulatory in nature, the inactive dynein/dynactin complexes must rely on a different interaction with TRAK to maintain association when the DRH is in the “closed” conformation. We tested whether the Spindly motif, a conserved dynactin binding site on TRAK, is required for inactive dynein/dynactin association using a deletion mutant. Both CC1-box and Spindly motif mutations reversed the net transport of TRAK2-driven cargo from predominantly dynein-driven to predominantly kinesin-driven (fig. S6D). However, compared to only a modest effect of the CC1-box mutant on dynein/dynactin association, the Spindly motif mutant eliminated most dynein/dynactin binding to both TRAK1 and TRAK2 (Fig. 2I, fig. S6E). These results indicate that the TRAK Spindly motif is required for dynein/dynactin association, and the additional binding of the LIC to the CC1-box is needed for activation of motility.

### Stress-induced phosphorylation of TRAK2 activates dynein

The dynein activating effect of the DRH-disrupting mutants raised the question of whether the association of the DRH with the CC1-box might be subject to physiological regulation. To explore this possibility, we used mass spectrometry to identify post-translational modification sites on TRAK co-immunoprecipitated with our synthetic cargo (fig. S7 A-D). Notable among the post-translationally modified sites was residue S84 that immediately precedes the DRH, and is conserved in TRAK2 (Fig. 3 A and B, fig. S7E). In TRAK1, a threonine is found at the equivalent site (T84), and this residue was not detected as phosphorylated in our mass spectrometry studies (fig. S7D).

**Fig. 3.**
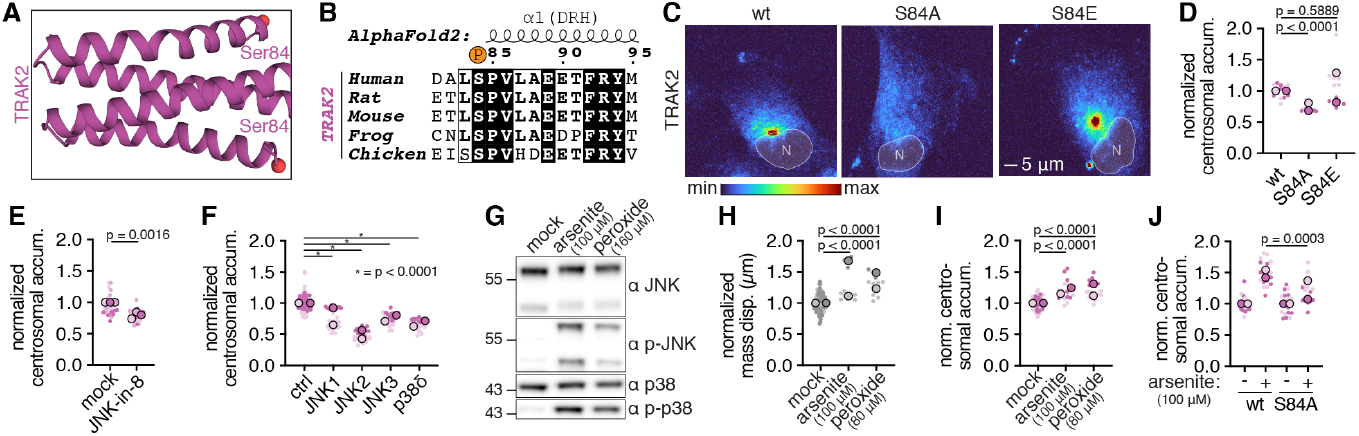
Activation of dynein by stress-induced phosphorylation of TRAK2. Representative maximum intensity projection images corresponding to panels E, F, H, I, J are shown in fig. S8 A, C, D, E, F respectively. (**A**) The TRAK2 S84 phosphorylation site identified by mass spectrometry highlighted in an inset showing the CC1-box region of the TRAK2 coiled-coil prediction as in fig. S4A (top). (**B**) S84 conservation in TRAK2. (**C**) Representative maximum intensity projection images of cargo from cell lines expressing wt, S84A or S84E TRAK2. (**D**) Quantification of perinuclear cargo accumulation in cells as shown in (C) from two biological replicates. n(wt)=12; n(S84A)=12; p<0.0001 (two-tailed t-test, t=6.615, df=22); n(S84E)=12; p=0.5889 (two-tailed t-test, t=0.5485, df=22). (**E**) Quantification of perinuclear cargo accumulation in cells expressing synthetic cargo and TRAK2 following mock or JNK-in-8 (10 μM) treatment (3h) from three biological replicates. n(mock)=27; n(JNK-in-8)=27; p<0.0016 (two-tailed t-test, t=3.332, df=52). (**F**) Quantification of perinuclear cargo accumulation in cells expressing TRAK2 following RNAi treatment as indicated from two biological replicates. n(wt)=120; n(JNK1,p38δ)=29, n(JNK2,JNK3)=30; one-way ANOVA (F= 4.271, dFn =4, dFd =233) with a post hoc Dunnett’s test was used to determine significance. (**G**) Immunoblot analysis of untransduced whole cell lysate showing JNK- and p38-kinase activation in response to 3h oxidative treatments as indicated. (**H**) Quantification of mitochondrial clustering following oxidative treatment in untransduced cells from two biological replicates. n(mock)=63; n(arsenite)=18; p<0.0001 (two-tailed t-test, t=6.356, df=113); n(peroxide)=13; p<0.0001 (two-tailed t-test, t=7.832, df=113). (**I**) Quantification of perinuclear cargo accumulation following oxidative treatment in cells expressing TRAK2 from two biological replicates. n(mock)=102; n(arsenite)=13; p<0.0001 (two-tailed t-test, t=6.436, df=79); n(peroxide)=13; p<0.0001 (two-tailed t-test, t=7.336, df=79). (**J**) Quantification of perinuclear cargo accumulation following arsenite treatment in cells expressing wt or S84A TRAK2 from two biological replicates. Arsenite-induced accumulation was normalized to mock treated cells for both variants. n(treated wt)=18; n(treated S84A)=18; p=0.0003 (two-tailed t-test, t=3.996, df=34).

To examine the role of phosphorylation of S84 in TRAK2-mediated transport, we generated a S84A mutant to block phosphorylation, and the negatively-charged S84E mutant as a potential phosphomimetic. The S84A mutant reduced TRAK2 dynein transport towards the centrosome compared with wildtype TRAK2 and the S84E mutant (Fig. 3 C and D). This result suggests that phosphorylation of S84 plays a role in regulating dynein transport, potentially by promoting the open conformation of the DRH.

Searching the sequence surrounding the TRAK2 S84 phosphorylation site against known consensus phosphorylation sites revealed that it best matched the proline-directed sites for stress-activated MAP kinases (SAPK) JNK1-3 and p38δ (*36*). To test whether this class of kinases is involved in activating TRAK2 dynein activity, we used both pharmacological and genetic perturbations. Treatment with the JNK kinase inhibitor JNK-in-8 (*37*) decreased TRAK2-mediated retrograde transport (Fig. 3E, fig. S8A). Individual RNAi knockdowns of several stress-activated kinases also diminished dynein transport, with JNK2 producing the strongest effect (Fig. 3F, fig. S8 B and C).

We next tested whether placing hTERT RPE-1 cells under conditions of oxidative stress led to activation of dynein motility. Arsenite and peroxide, both inducers of oxidative stress, activated JNK and p38 MAP kinase in these cells (Fig. 3G). We also found that these agents caused perinuclear clustering of mitochondria (*29, 38*) (Fig. 3H, fig. S8D), as well as TRAK2-driven synthetic cargo (Fig. 3I, fig. S8E). We next explored whether the phenomena of stress-activated MAP kinase activation and increased dynein motility might be causally linked through stress-activated phosphorylation of TRAK S84. Indeed, the TRAK2 S84A mutant showed attenuated dynein activation in response to oxidative stress compared with wild type TRAK2 (Fig. 3J, fig. S8F).

### The DRH determines transport direction

Prompted by our finding of different stress-induced phosphorylation of TRAK1 and 2, we asked whether the overall distinct directional transport preferences of these two adaptor proteins are conferred by their DRH and adjacent phosphorylation site (aa 81-106), or if they require other TRAK domains. To test this, we generated chimeric proteins with residues 81-106 swapped between TRAK1 and TRAK2 (Fig. 4 A-D). The results clearly show that the directional preference was determined by the source (TRAK1 or TRAK2) of the DRH in the chimera. Since the DRH only constitutes ∼3% of the TRAK polypeptide, this result indicates that the DRH region plays an important role in tuning the motor preference of TRAK between anterograde kinesin and retrograde dynein transport.

**Fig. 4.**
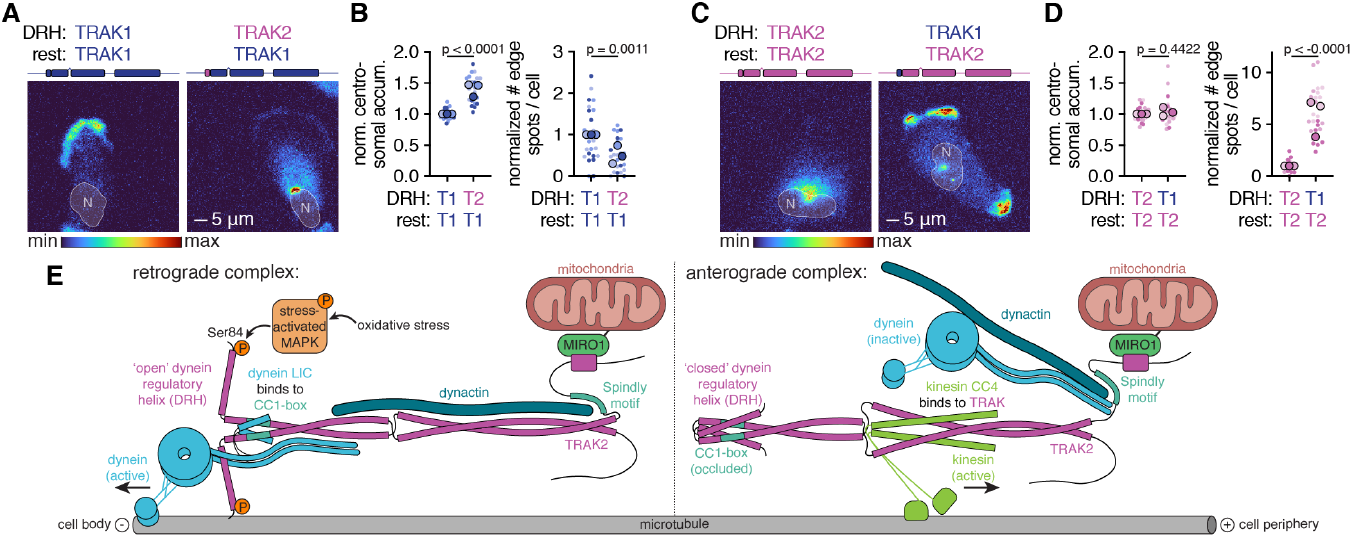
The DRH determines transport direction. (**A**) Representative maximum intensity projection images of cargo from cell lines expressing TRAK1 containing either the endogenous (left) or TRAK2 (right) DRH sequence. (**B**) Phenotype quantification from three biological replicates as shown in (A). For perinuclear accumulation n(TRAK1 DRH)=27; n(TRAK2 DRH)=27; p<0.0001 (two-tailed t-test, t=9.838, df=52). For peripheral accumulation n(TRAK1 DRH)=26; n(TRAK2 DRH)=27; p=0.0011 (two-tailed t-test, t=3.457, df=51). (**C**) Representative maximum intensity projection images of cargo from cell lines expressing TRAK2 containing either the endogenous (left) or TRAK1 (right) DRH sequence. (**D**) Phenotype quantification from three biological replicates as shown in (C). For perinuclear accumulation n(TRAK2 DRH)=27; n(TRAK1 DRH)=26; p=0.4422 (two-tailed t-test, t=0.7745, df=51). For peripheral accumulation n(TRAK2 DRH)=27; n(TRAK1 DRH)=25; p<0.0001 (two-tailed t-test, t=10.91, df=50). (**E**) Model for directional switching of mitochondria dictated by the TRAK DRH conformation (see main text for more details).

## DISCUSSION

In summary, we find that the coordination of dynein and kinesin motors on mitochondria is determined by the conformational state of the TRAK dynein regulatory helix (DRH) which modulates the accessibility of the dynein LIC to the TRAK CC1-box (Fig. 4E). In the anterograde TRAK conformation, the DRH is closed (folded-back) such that the dynein LIC cannot bind the TRAK CC1-box; this conformation allows kinesin to bind the TRAK coiled-coil to mediate anterograde transport. The anterograde TRAK conformation remains associated with dynein/dynactin through its Spindly motif, allowing kinesin to carry dynein/dynactin as an inactive cargo, as was also observed in in vitro motility assays (*24*). In the retrograde TRAK conformation, the DRH is open, allowing the dynein LIC to engage with the TRAK CC1-box; the dynein LIC interaction with the CC1-box may be necessary for aligning the dynein motor domains in a parallel manner to engage in processive movement (*39*). This active dynein conformation is incompatible with, and displaces bound kinesin from TRAK, mitigating interference from the opposing motor. Since the DRH is distal from the kinesin binding site on TRAK, the structural mechanism for this displacement is not known. The active conformation of dynein/dynactin could either sterically clash with kinesin, or stabilize the TRAK coiled-coil such that it can no longer open up to form the four-helix bundle with the kinesin heavy chain.

Our results also provide potential insight into, and raise new questions about the regulation of directed mitochondrial transport. TRAK1 and TRAK2 appear to be differentially tuned by the closed (anterograde) or open (retrograde) conformation of their DRHs. This differential tuning may help govern the overall cellular distribution of mitochondria, along with mechanisms that regulate the attachment of mitochondria to the actin cytoskeleton (*31*). The two TRAKs may also control mitochondrial distribution in response to different intracellular cues. TRAK2, which is activated by stress-activated MAP kinases, is enriched in the brain(*11*), which is particularly sensitive to ROS generation (*40*). The DRH of TRAK1 appears to be primarily in a closed state in our cell culture system, favouring kinesin transport. However, TRAK1 contains a fully functional CC1-box and can be artificially activated for dynein transport by mutagenesis. Thus, TRAK1 may activate dynein transport by DRH opening in response to other, yet unidentified, physiological signals. Regulation of dynein LIC binding to TRAK by the mechanisms identified in this work may be shared among other bidirectional adaptor proteins, such as lysosome-transporting JIP3 (autoinhibitory helix (*41*)) or endosome-transporting HOOK3 (association with inactive dynein/dynactin (*42, 43*)), although the signalling networks that tune these features in other adaptors have not yet been identified. Our synthetic cargo transport assay, which could in principle be applied to these and other motor adaptors, should prove useful for dissecting future questions concerning microtubule motor regulation in cells.

## NOTES AND REFERENCES

## Acknowledgments

We thank N. Stuurman, M. DeSantis and D. Alcor for assistance with image acquisition, K. Schaefer and C. Li for their assistance with flow cytometry, L. Lavis for generously providing reagents, members of the Vale lab for reagents and discussion, J. Lippincott-Schwartz and members of her lab for discussion, A. S. Moore for advice on image presentation. We are grateful to H. Farrants, C. Ott, J. Sheu-Gruttadauria and T.E.T. Mevissen for critical feedback on the manuscript.

## Funding

Howard Hughes Medical Institute, National Institute of General Medical Sciences grant R35GM147554 (SAM), HHMI fellowship of The Jane Coffin Childs Memorial Fund for Medical Research (CG), EMBO Long-term fellowship grant ALTF 99-2019 (CG)

## Author contributions

Conceptualization: CG, RDV; Formal analysis: CG, WPG, SAM; Software: WPG; Funding acquisition: RDV, CG; Investigation: CG, MGPS, SAM; Methodology: CG, YW; Project administration: CG, RDV; Resources: YW; Supervision: CG, RDV; Visualization: CG, SAM; Writing – original draft: CG, RDV; Writing – review & editing: CG, MGPS, WPG, SAM, YW, RDV

## Competing interests

Authors declare that they have no competing interests.

## Data and materials availability

All data are available in the main text or the supplementary materials. All reagents and materials are available from C.G. upon reasonable request. All generated code is available at Zenodo (https://doi.org/10.5281/zenodo.13150980).

## MATERIALS AND METHODS

### Molecular Biology

Miro1-sfGFP-FLAG-I3-01, Miro1-Halo-FLAG-I3-01 and TRAK-mScarlet and fusion constructs were assembled using the NEBuilder HiFi DNA Assembly Master Mix (New England Biolabs) into a 2^nd^ generation lentiviral transfer vector containing a SFFV promoter to drive expression as well as cPPT and WPR elements to enhance transduction and expression respectively (gift from Kara L. McKinley). Human Miro1, TRAK1 and TRAK2 sequences were obtained either as cDNA, or gBlocks from IDT. I3-01 (*32*), sfGFP, mScarlet, Halo and FLAG sequences were amplified from laboratory vector stocks. A monomerizing V206R mutation was introduced in the sfGFP sequence. The Miro1 fusion plasmids contain Miro1 truncated before the transmembrane domain (aa 591-618), while TRAK1 and TRAK2 constructs are truncated before their mitochondrial binding domain (before aa 703-953 and aa 701-914 respectively). Site-directed mutagenesis primers were designed using the QuikChange protocol, and used to amplify two overlapping mutagenized fragments of the insert of interest. Fragments were then assembled back into the original vector using NEBuilder HiFi DNA Assembly Master Mix (New England Biolabs).

### Cell culture, cell line generation and perturbation

hTERT RPE-1 (cTT33.1 carrying inducible Cas9, gift from Kara L. McKinley and Iain Cheeseman) and Lenti-X HEK293T cells for lentiviral packaging (Takara Bio) were cultured in complete media: high-glucose DMEM (Gibco) supplemented with 10% tetracycline negative FBS (Corning) and 1x Penicillin-Streptomycin-L-Glutamine (Corning). For passage, cells were dissociated using 0.05% Trypsin/0.53mM EDTA (Corning) following a PBS rinse. Low cell passage number was maintained, and cells were checked regularly for Mycoplasma using the MycoAlert PLUS Mycoplasma Detection kit (Lonza).

#### Stable cell line generation

Stable cell lines were generated by lentiviral transduction of the parent hTERT RPE-1 clone, and the transduced cell population was further enriched by FACS. A polyclonal population was used for all experiments, all comparisons are drawn between cells handled (transduced, gated and grown) identically. Briefly, lentivirus was generated by transfecting 1×10^6 Lenti-X HEK293T cells with transfer vector (2 μg), packaging vector (1 μg) and VSV-G envelope vector (0.5 μg) using Lipofectamine LTX reagent (Thermo Fisher) and Opti-MEM I Reduced Serum Medium (Gibco). Transfection-mix containing media was exchanged the following day, and crude viral supernatant was harvested 72 hours after transfection. Obtained viral supernatant for Miro1-FP-FLAG-I3-01 and TRAK-mScarlet fusions (20 μL) was mixed with hTERT RPE-1 cells (0.05×10^6) in suspension containing polybrene at 5 μg/mL. The resulting mixture was centrifuged in a culture plate at 1000xg for 1h in 24-well tissue culture dishes (estimated MOI 100). Once confluent, cells were expanded for FACS. Gates were set based on untransduced or hTERT RPE-1 cells transduced with virus containing a single FP to select for low/medium expressors. See Supplementary Fig. 1 for representative gating strategy.

#### Gene knockdown using siRNA

Silencer Select (JNK1 - s11152, JNK2 - s229708, JNK3 - s11161, p38δ - s11164), Validated Silencer Select (KIF5B - s731, DYNC1H1 - s4202) and the Silencer Select Negative control #1 siRNAs were ordered from Thermo Fisher. siRNAs were reverse transfected at a final concentration of 25nM using Lipofectamine RNAiMAX (Thermo Fisher) and Opti-MEM I Reduced Serum Medium (Gibco). For imaging, 0.5×10^4 hTERT RPE-1 cells were seeded into 96 square-well glass-bottom imaging dishes (Matriplate MGB096-1-2-LG-L). For WCL analysis, 2.5×10^4 hTERT RPE-1 cells were seeded into 12-well tissue culture dishes. Cells were imaged, fixed or harvested 72 hours after siRNA transfection.

#### Drug treatments

As above, 96- and 12-well dishes were used for imaging and WCL analysis respectively. Drug concentration and treatment duration is indicated in the figures and legends where applicable. A 2x solution of nocodazole, CCCP, arsenite and tert-Butyl hydroperoxide (Sigma-Aldrich), or JNK-in-8 (MedChemExpress) was prepared in complete culture media, which was applied at a 1:1 ratio with pre-existing culture media. Mock treatment consisted of addition of media alone. For imaging, cells were either fixed at the treatment endpoint (CCCP, Arsenite, peroxide) or imaged live at the treatment endpoint (nocodazole, JNK-in-8). For whole cell lysate immunoblot analysis, cells were harvested at the treatment endpoint.

### Image acquisition - Synthetic cargo accumulation, mitochondrial network

#### Sample preparation

Cells were seeded into 96 square-well glass-bottom imaging dishes (Matriplate MGB096-1-2-LG-L), such that they reach a density of ∼4×10^4 cells/well at the time of imaging, and at least a day prior to imaging. For assessing the effects of TRAK variants, cells were stained with Hoechst 33342 (Thermo Fisher) prior to live imaging. Where indicated, cells were also stained with MitoTracker DeepRed FM (Thermo Fisher). For most cell perturbation experiments, cells were fixed using 4% PFA (Electron Microscopy Sciences) diluted in PHEM buffer (60 mM PIPES (from pH = 6.8 stock), 25 mM HEPES (from pH = 7.4 stock), 10 mM EGTA, 5 mM MgCl2), rinsed with PBS and stained with Hoechst 33342 prior to imaging. To assess perturbation effects on the mitochondrial network, cells were fixed as above, permeabilized using 0.5% Triton-X in PHEM, blocked using 3% BSA in PBS, incubated overnight at 4°C with primary antibody (Tom20; BD Biosciences, 612278 or Tom20; abcam, ab186734), washed, incubated with AlexaFluor-labelled secondary antibody (Thermo Fisher, A-11001 or A-21244) and Hoechst 33342.

#### Image acquisition

All images were acquired on an Eclipse Ti2 inverted microscope (Nikon) equipped with a CSU-W1 spinning disk confocal unit (Yokogawa) using a 50 μm-pinholed disk, and controlled via NIS-Elements (Nikon, AR 5.42.03). The system was equipped with Hamamatsu ORCA-Fusion BT sCMOS cameras (Hamamatsu) and operated in ‘ultra-quiet’ mode with an exposure time of 200 ms. The environment was maintained at 37°C with 5% CO_2_ using either a cage or a stage top incubation system (Okolab). Cells were excited with 405-nm (nuclei), 488-nm (synthetic cargo, Tom20 immunofluorescence), 561-nm (TRAK), and 640-nm (MitoTracker DeepRed, Tom20 immunofluorescence) laser lines and emission was collected using one of the following objectives (Nikon) and standard filter sets: 20x/0.75 NA Plan Apo λ or 20x/0.8 NA Plan Apo λ D (high content acquisition), 60x/1.4 NA Plan Apo λ D (Fig. 1c), 100x/1.45 NA Plan Apo λ D (Extended Data Fig. 2a, Video 1). For typical high content acquisition using the 20x objectives, the JOBS module was utilized and 16 μm z-stacks were acquired using a piezo-driven stage with a 2 μm step size (Mad City Labs).

### Image acquisition - Synthetic cargo particle tracking

#### Sample preparation

Cells were seeded into 96 well glass-bottom imaging dishes as above. Prior to imaging, cells were stained with 100 nM PA-JF646-Halo photoactivatable dye for 30 min in complete media (gift from L. Lavis, Janelia Research Campus (*33*)) and washed 3x for 10 min with complete media.

#### Image acquisition

Images were acquired on the same spinning disk confocal microscope described above using a 50 μm-pinholed disk operated in ‘ultra-quiet’ mode with and exposure time of 200 ms/frame. Prior to photoactivation, cells were either imaged using transmitted light (no TRAK expression) or the 561 nm laser (TRAK1 or TRAK2 expression) to infer the cellular footprint and nuclear position necessary to determine position of photoactivation and to enable computational track reorientation (see analysis below). For simultaneous imaging and dye activation, live acquisition was started before stimulating cells using a 405 nm laser controlled via an Opti-Microscan for photostimulation module targeting a circular ROI (1.5 μm diameter) halfway between the nucleus and cell periphery. Emission was collected using a 100x/1.45 NA Plan Apo λ objective (Nikon) using a 705/72 nm filter.

### Image analysis

#### Synthetic cargo accumulation, mitochondrial network

A Cell Profiler (*44*) (4.2.4) pipeline was developed for analysis of high content imaging of synthetic cargo accumulation (see Extended Data Fig. 3). Nuclei (30-80 px diameter) were segmented from the hoechst channel using global otsu thresholding, the threshold correction factor was adjusted (0.5-1.1) for best performance across the analyzed image set. To measure perinuclear accumulation of the synthetic cargo, the segmented nuclear mask was expanded by 10 px to define the perinuclear region. A custom plugin was developed to measure the degree of pixel intensity inequality within the perinuclear region mask of the synthetic cargo image by calculating the Gini coefficient (https://doi.org/10.5281/zenodo.13150980). The Gini coefficient takes values from 0-1, and compares the observed pixel frequency distribution to a hypothetical case where all pixels have a uniform intensity value. To identify edge spots, the segmented nuclear mask was expanded by 15 px and subtracted from the synthetic cargo image such that only pixels corresponding to peripheral cell areas remained. Bright spots (>6 standard deviations above background, min 5px diameter) were identified in the resulting image using global robust background thresholding. Values >2 STDEV above the mean were excluded, as they are likely to result from segmentation error. A custom script was used to process the Cell Profiler output (https://doi.org/10.5281/zenodo.13150980). Briefly, we took the Gini values calculated for each cell and took the median over all cells in each field of view. We also normalized the number of edge spots observed in each field of view by the number of detected nuclei. These per-field-of-view values were then plotted as a single datapoint for each condition.

To measure the degree of mitochondrial clustering, the Cell Profiler and post-processing pipelines above were modified to measure the intensity mass displacement of the mitochondrial image within the perinuclear region mask. The mass displacement corresponds to the distance (in μm) between the centres of gravity in the grey-level representation of the object and the binary representation of the object.

#### Synthetic cargo particle tracking

Acquired time-lapse series were denoised using the Denoise.ai function of the NIS.ai package (Nikon) to remove shot noise. Particle trajectories were generated using the TrackMate Fiji plugin (*45*). To reduce linking artifacts caused by high particle density immediately following photoactivation, only frames acquired 40-120s (200ms per frame, 400 frames) after photoactivation were included in the analysis. A Laplacian of Gaussian detector was used to identify synthetic cargo particles with an estimated object diameter of 0.4 μm, employing sub-pixel localization. The linear assignment problem tracker was used to generate trajectories with a max linking distance and a gap closing distance set to 1.0 μm. Only tracks containing 6 or more spots were included in the analysis to reduce the effect of out-of-plane diffusion. For each cell, coordinates of the bleach spot and the approximate nucleus centroid were recorded from a bright-field or TRAK channel image and used to generate a cell orientation vector between the nucleus to the bleach spot (periphery). Kymographs in Fig. 1e were generated in ImageJ by re-slicing acquired image stacks using a rectangular ROI (8×0.65 μm) as indicated in Extended Data Fig. 2b.

Trajectories acquired for cells expressing no TRAK (14 cells, 3,789), TRAK1 (21 cells, 24,392), TRAK2 (47 cells, 32,669) across three different days were exported from TrackMate and analysed using a custom script (https://doi.org/10.5281/zenodo.13150980). Briefly, all tracks were origin aligned and reoriented according to the experimentally determined cell orientation vector, such that the cell periphery is in the +ve y direction and the nucleus in the -ve y direction. By comparing the properties of particle tracks obtained with and without TRAK expression, we defined tracks with >3.0 μm maximum distance travelled as processive (no TRAK: 8, TRAK1: 573, TRAK2: 270). To assess the distribution of track properties, the final position of origin-aligned reoriented tracks, the mean instantaneous speed per track and the mean instantaneous speed per track per final position was plotted for all processive tracks in the TRAK1- and TRAK2-expressing condition.

### Biochemistry

#### Immunoprecipitation

For each replicate, cells from a confluent 15-cm dish were washed with, and scraped into ice-cold PBS supplemented with cOmplete Protease Inhibitor Cocktail, PhosSTOP (Roche) and 1 μM Thiamet G (Tocris), centrifuged at 200xg, and frozen at -80°C. FLAG immunoprecipitation protocol was based on the resin manufacturers recommendation (ChromoTek DYKDDDDK Fab-Trap™ Agarose, Proteintech). Cells were lysed in IP buffer (25 mM HEPES (from pH = 7.4 stock), 50 mM KOAc, 2 mM MgOAc, 1 mM EGTA, 1% v/v glycerol, 1x cOmplete Protease Inhibitor Cocktail (Roche), 1x PhosSTOP (Roche), 1 μM Thiamet G (Tocris), 0.5 mM TCEP (Sigma-Aldrich), 0.5 MgATP (Thermo Fisher), 1 mM PMSF (Calbiochem)) supplemented with 0.2% Triton X-100. Lysate was clarified by centrifugation at 17,000xg and diluted 2:3 with IP buffer. Diluted lysate was applied to DYKDDDDK Fab-Trap Agarose resin equilibrated with IP buffer, resin washed 3x with IP buffer, and protein eluted directly in SDS-sample buffer by boiling resin for 5 min at 95°C.

#### WCL preparation

Cells grown on a 12-well tissue culture dish were washed with ice-cold PBS. Cells were scraped into RIPA lysis buffer (Thermo Fisher) supplemented with 1x cOmplete Protease Inhibitor Cocktail (Roche), 1x PhosSTOP (Roche), 1 μM Thiamet G (Tocris), 0.5 mM TCEP (Sigma-Aldrich), 0.5 MgATP (Thermo Fisher), 1 mM PMSF (Calbiochem). Lysates were sonicated using at 20% power (Q125 Sonicator, QSonic) for a total of 20s (5s on, 5s off). Total protein concentration was measured using a BCA Protein Assay (Thermo Fisher) and equalized across all samples analyzed in each experiment.

#### Western blotting

Immunoprecipitated or whole cell lysate samples were resolved on 4-12% Bis-Tris SDS NuPAGE gradient gels (Invitrogen) together with Broad Protein Standard (New England Biolabs), and transferred to a nitrocellulose membrane using the Trans-Blot Turbo Transfer system (Mixed MW transfer program, BioRad). For resolving the dynein heavy chain (DHC1, 532 kDa), a 3-8% Tris-Acetate SDS NuPAGE gradient gels (Invitrogen) was used together with the High MW transfer program instead. Membranes were blocked in a 5% (w/v) milk solution (Fisher Sci) in PBS-T (PBS + 0.1% (v/v) Tween-20) and incubated overnight at 4°C with primary antibody diluted in PBS-T + 3% (w/v) BSA + 0.02% (w/v) NaAzide (see below for a list of primary antibodies used). The membranes were then washed with PBS-T and incubated overnight at 4°C with secondary antibody diluted in a 5% (w/v) milk solution in PBS-T (HRP-conjugated Goat anti-Mouse or Goat anti-Rabbit Secondary Antibody, Thermo Fisher), washed with PBS-T and visualized using the SuperSignal West Pico PLUS Chemiluminescent Substrate and the ChemiDocTouch Imaging System (BioRad).

The following primary antibodies were used: GFP (CST, 2555S), RFP (Proteintech, 6g6), KIF5B (abcam, ab167429), DHC1 (Santa Cruz Biotechnology, sc-514579), p150 (BD Biosciences, 610473), GAPDH (CST, 2118S), DIC (Sigma-Aldrich, MAB1618), p50 (BD Biosciences, 611002), SAPK/JNK (CST, 9252S), Phospho-SAPK/JNK (Thr183/Tyr185) (CST, 9251S), p38 MAPK (CST, 9212S), Phospho-p38 MAPK (Thr180/Tyr182) (CST, 9211S), O-GlcNAc [RL2] (abcam, ab2739), p38δ MAPK (CST, 2308T).

For quantification of immunoprecipitation experiments, western blot band intensity from signal corresponding to the GFP-labelled synthetic cargo and motor protein components of interest as indicated (KIF5B, DHC1, p150) was quantified in ImageJ. The band intensity corresponding to co-eluting motor protein components was normalized by the intensity of the immunoprecipitated synthetic cargo (GFP band), and subsequently normalized with respect to a chosen condition as indicated to allow comparison between biological replicates.

### Mass Spectrometry

To obtain sufficient protein for band excision from a Coomassie-stained protein gel, the immunoprecipitation protocol described above was scaled up eight-fold. An aliquot of the eluted SDS sample was used for validation of the experiment by IP-western (Extended Data Fig. 7a), and the remainder of the sample was resolved on a 4-12% Bis-Tris SDS WedgeWell Bolt gradient gel (Invitrogen), stained with InstantBlue Protein Stain (Expedeon) and bands excised as indicated in Figure (Extended Data Fig. 7b).

Gel bands were digested with trypsin (Promega), and desalted as previously described (*46*). Data was acquired on a Thermo Orbitrap Eclipse with ETD. Peptides were separated on a 60 min effective gradient from 2% solvent B (80% acetonitrile, 0.1% formic acid in water) to 30% B. The MS1 scan was measured in the Orbitrap at 60,000 resolution with a 50 ms injection time for 40,000 charges. The scan range was 350-1800 m/z for 2-8 plus precursors. MS2 scans were performed after a 2.0 m/z isolation window by the quadrupole, filled to 50,000 charges in 105 ms, followed by HCD at a normalized collision energy of 28%. Product ions were read out in the Orbitrap at 15,000 resolution. If 204.0867 or 138.0545 m/z +/- 15 ppm was detected in the MS2 scan, the mass spectrometer refilled for up to 250 ms for 100,000 charges before performing EThcD at 25% supplemental activation. Three microscans were permitted and product ions were measured in the Orbitrap at 7,500 resolution. Data was searched using Byonic (Protein Metrics, Inc) for protein identifications. Only proteins identified in the sample were further searched for HexNAc (STN), common 3, phosphorylation (STY) common 2. Cys carbamidomethylation was a fixed modification and protein N-term acetyl and Met oxidation were rare. All discussed site assignments were validated by hand.

### AlphaFold2 prediction and TRAK sequence analysis

Structure predictions were performed using AlphaFold2-Multimer (*47*) implemented via ColabFold running on the Colab platform (Google). Two copies of TRAK1 (aa 1-396) and TRAK2 (aa 1-914) were used to generate predictions N-terminal coiled-coil predictions shown in Fig. 2a (top) and Extended Data Fig. 4a (top). Two copies of TRAK1 or TRAK2 (aa 1-360) were used together with two copies of KIF5B (aa 821-963) to generate kinesin complex predictions shown in Fig. 2a (bottom) and Extended Data Fig. 4a (bottom). Structural figures were generated using PyMol (www.pymol.org). pLDDT scores generated by AlphaFold2-Multimer were mapped onto the structure using indicated LUTs (Pymol LUT implementation designed by Konstantin Korotkov). Sequence alignment with structure prediction mapping in Extended Data Fig. 1 was generated using ESPript (*48*).

### Statistics and reproducibility

Statistical analysis and graphing were carried out using Prism (v 9.2.0). Superplots representing imaging data show means for each biological replicate superimposed on the calculated phenotypic scores for individual fields of view from samples handled in parallel (see Extended Data Fig. 3). Sample number, the number of biological replicates and employed statistical tests are described in each figure legend. Two tailed unpaired t-tests were used to determine significance when one or two experimental groups were compared to a control distribution, otherwise ANOVA with post hoc Dunnett’s testing was used. The TRAK1 control distribution is identical in the comparisons quantified in Fig. 2d, 2g, 4b, as the experiments were carried out in parallel (transduction, FACS, imaging). The TRAK2 control distribution is identical in the comparisons quantified in Extended Data Fig. 4f, 6b, 6d and Fig. 4d, as the experiments were also carried out in parallel (transduction, FACS, imaging). All representative micrographs were selected from matching cell lines handled in parallel.

### Data availability

The datasets generated during and analysed during the current study are available from C.G. upon reasonable request.

### Code availability

Custom code was essential to the conclusions of this work. All generated code is available at Zenodo (https://doi.org/10.5281/zenodo.13150980).

---

**Fig. S1.**
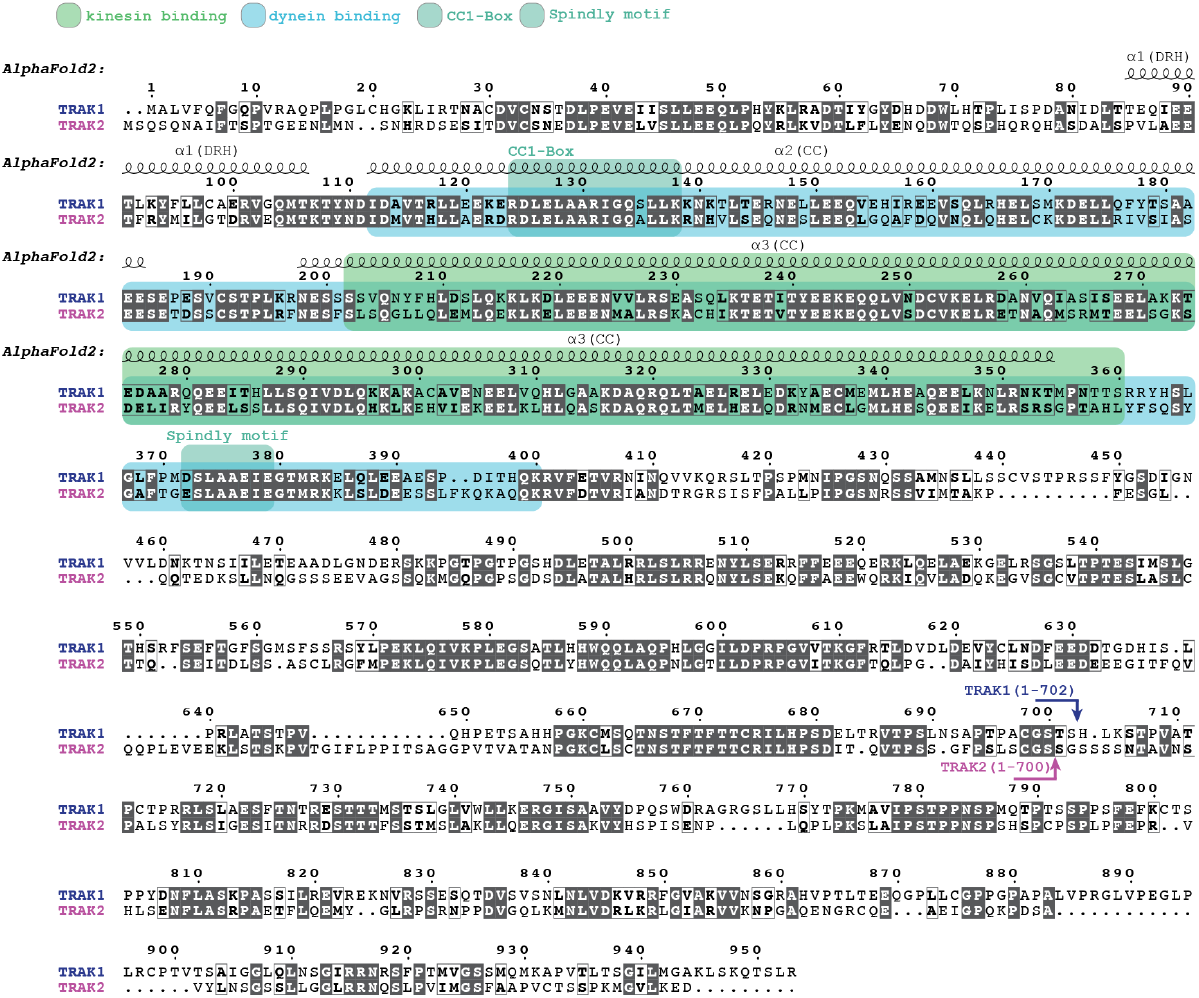
Sequence alignment of TRAK1 and TRAK2. Sequence alignment of TRAK1 and TRAK2 with the following annotations: previously mapped binding regions for kinesin (green) and dynein (blue), CC1-Box and Spindly motif (cyan), secondary structure elements reliably identified by AlphaFold2 (see fig. S5), C-terminal boundaries for the constructs used in this study.

**Fig. S2.**
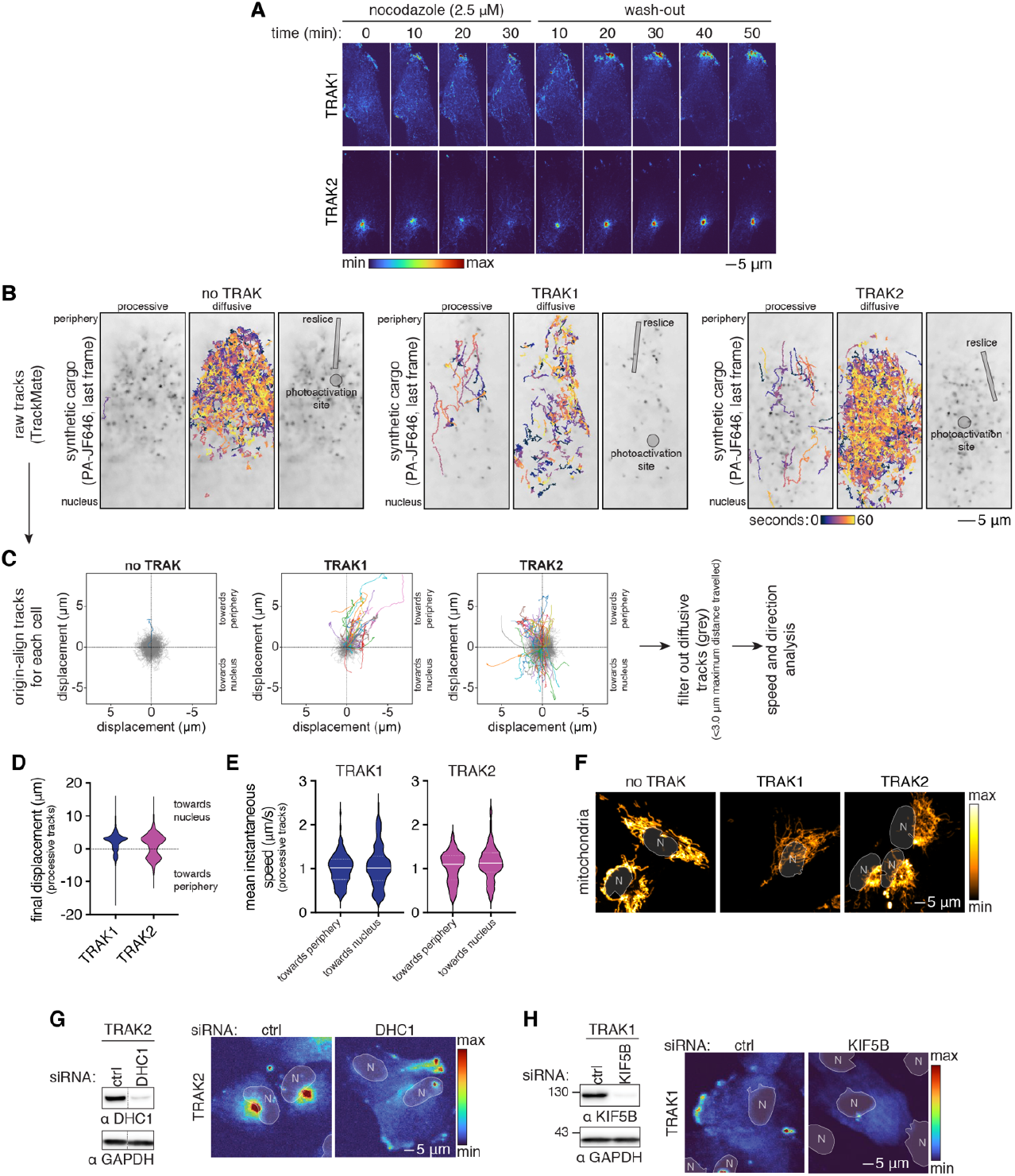
Synthetic mitochondrial cargo characterization. (**A**) Maximum intensity projections of frames from time-lapse imaging of cargo localization following nocodazole treatment (2.5 μM, 30 min) and wash-out (see Movie S1). (**B**) Final frames from time-lapse imaging of photoactivated synthetic cargo of cells expressing no TRAK, TRAK1 or TRAK2 (see Movie S2). Overlaid are processive (left panel, >3.0 μm maximum distance travelled) or diffusive (middle panel, <3.0 μm maximum distance travelled) trajectories generated by TrackMate for the previous 60s. Photoactivation site and source area for kymographs in Fig. 1E are overlaid on the right panel. (**C**) Origin-aligned tracks from cells as in (B) that have been oriented such that the cell periphery is in the +ve y direction and the nucleus in the -ve y direction. Processive tracks are highlighted in colour, diffusive tracks are grey. (**D**) The distribution of final particle positions from origin-aligned and reoriented processive tracks as in (C) from the TRAK1 (n = 21 cells, 573 tracks) and TRAK2 (n = 47 cells, 270 tracks) datasets. (**E**) Mean instantaneous speed averaged over the length of processive tracks (maximum distance travelled >3.0 μm) from the TRAK1 and TRAK2 datasets as above, separated by direction of movement. This speed was 1.00 ± 0.02 μm/s for TRAK1 trajectories oriented towards the periphery (n=460 tracks), 1.06 ± 0.04 μm/s for TRAK1 trajectories oriented towards the nucleus (n=113 tracks), 1.04 ± 0.03 μm/s for TRAK2 trajectories oriented towards the periphery (n=172 tracks), 1.13 ± 0.04 μm/s for TRAK2 trajectories oriented towards the nucleus (n=98 tracks). (**F**) Maximum intensity projection images showing the mitochondrial networks of cells (MitoTracker) displayed in Fig. 1C. The mitochondria of TRAK1 overexpressing cells are distributed similarly to those without TRAK expression, while the mitochondria of TRAK2 overexpressing cells are more clustered. (**G**) Left: Immunoblot analysis of whole cell lysate from cells expressing synthetic cargo and TRAK2 showing the efficacy of DHC1 siRNA treatments used in Fig. 1H. Right: Representative maximum intensity projection images of cargo following RNAi treatment as indicated. (**H**) Left: Immunoblot analysis of whole cell lysate from cells expressing synthetic cargo and TRAK1 showing the efficacy of KIF5B siRNA treatments used in Fig. 1I. Right: Representative maximum intensity projection images of cargo following RNAi treatment as indicated.

**Fig. S3.**
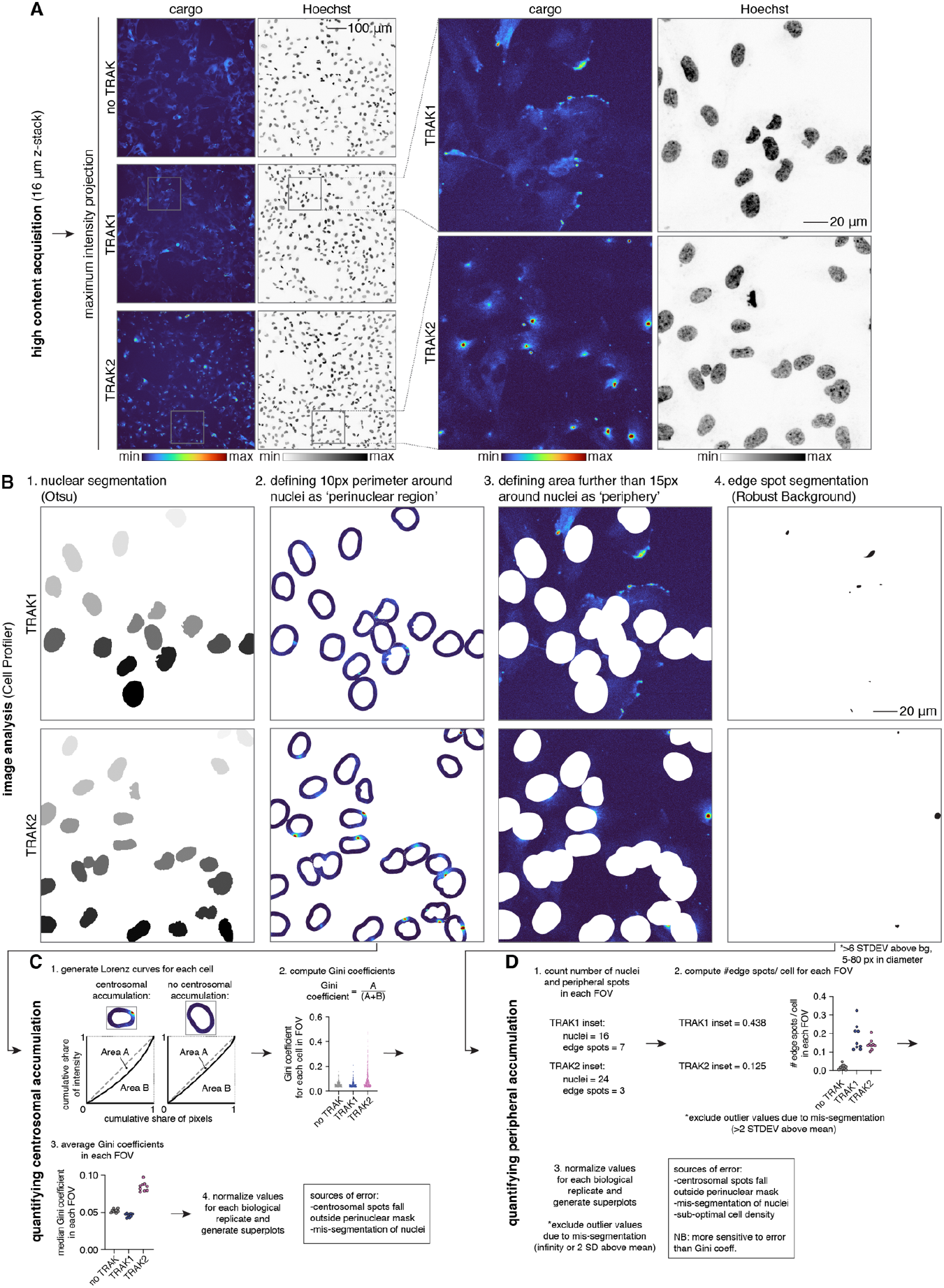
Quantifying synthetic cargo localization. (**A**) Left: Maximum intensity projection images showing an entire field of view (FOV) acquired for cells expressing synthetic cargo (left) and labelled with Hoechst (right) at 20x magnification. Right: Insets for TRAK1 and TRAK2 conditions showing O(10) cells. (**B**) Example outputs from the Cell Profiler pipeline used for phenotype quantification for the inset shown in (A). Nuclear segmentation (left) was used to mask the synthetic cargo image and define a ‘perinuclear region’ for each cell (middle left) and the ‘periphery’ for each FOV (middle right). Peripheral accumulations of cargo were identified by a subsequent round of segmentation (right). (**C**) A custom Cell Profiler plugin was used to calculate the Gini coefficient for the pixel intensity distribution in the perinuclear region of each segmented nucleus. The Gini coefficient takes values from 0-1 and measures the inequality of pixel intensities in each distribution by comparing the observed pixel frequency distribution to a hypothetical case where all pixels have a uniform intensity value (top). Median Gini coefficients are calculated for each FOV, normalized within each biological replicate and plotted (bottom). (**D**) The number of edge spots identified in the ‘peripheral’ segments of each FOV was normalized by the number of segmented nuclei. The number of edge spots per cell is calculated for each FOV, normalized within each biological replicate and plotted.

**Fig. S4.**
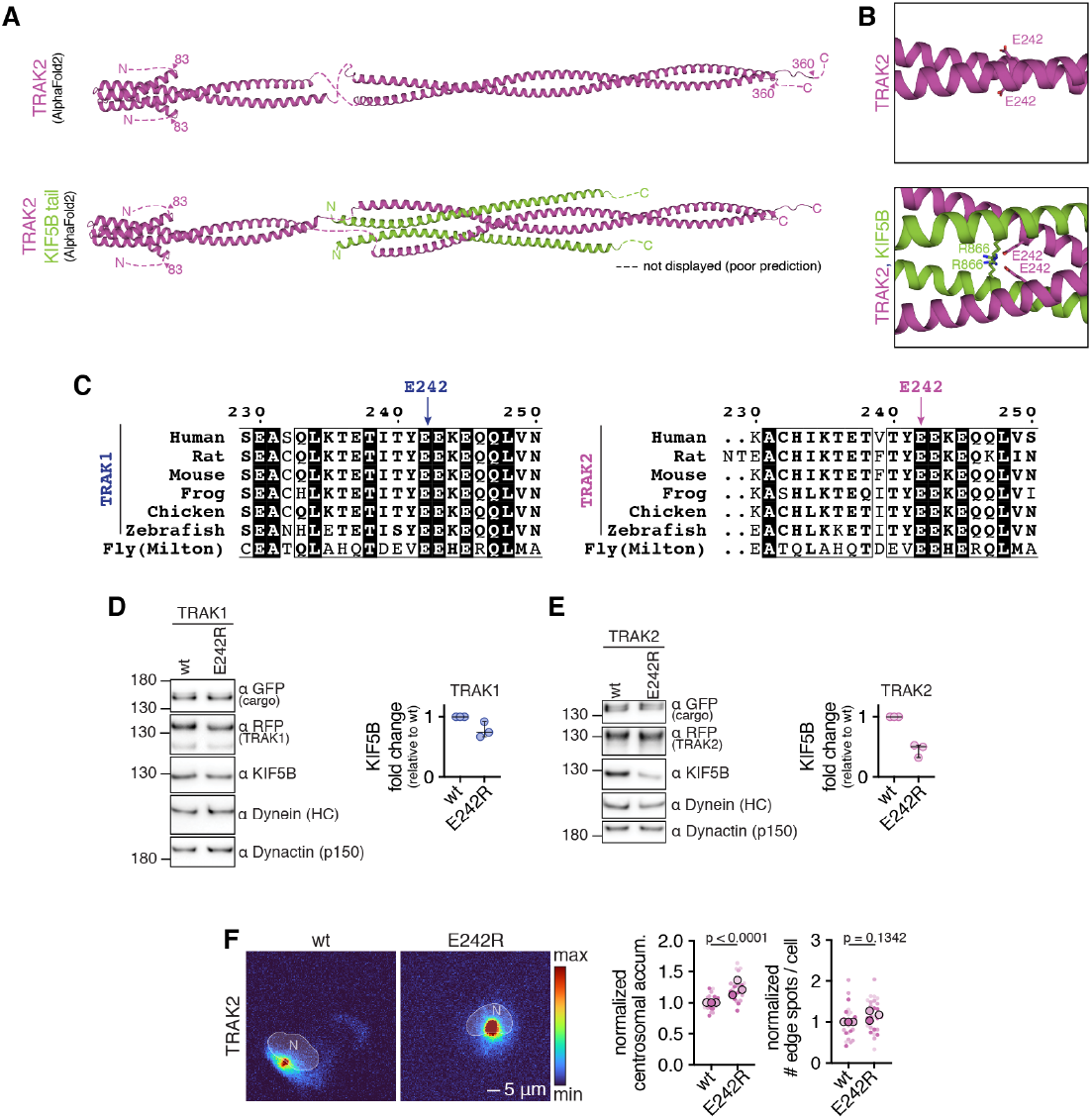
Probing the AlphaFold2 model for kinesin interaction. (**A**) AlphaFold2 predictions for TRAK2 (magenta). Top: N-terminal coiled-coil dimer (residues shown 83-360). Bottom: N-terminal coiled-coil dimer in complex with the CC4 region of kinesin-1 (green, residues shown 820-923). For clarity, the TRAK2 coiled-coil is split into two separately aligned sections at a site of disorder (see fig. S5). (**B**) Inset from the predictions in (A) showing residues E242 in TRAK1 and R866 in kinesin-1. (**C**) E242 conservation in TRAK1 and TRAK2. (**D, E**) Left: Immunoblot analysis of the motor-adaptor components associated with FLAG-purified cargo from indicated cell lines. Right: quantification of relative motor protein levels normalized to wt. Error bars show the median and 95% confidence interval; n = 2 for all conditions. (**F**) Left: Representative maximum intensity projection images of cargo from cell lines expressing wt or E242R TRAK2. Right: Phenotype quantification from three biological replicates as shown on the left. For perinuclear accumulation n(wt)=27; n(E242R)=27; p<0.0001 (two-tailed t-test, t=5.826, df=52). For peripheral accumulation n(wt)=26; n(E242A)=26; p=0.1342 (two-tailed t-test, t=1.523, df=50).

**Fig. S5.**
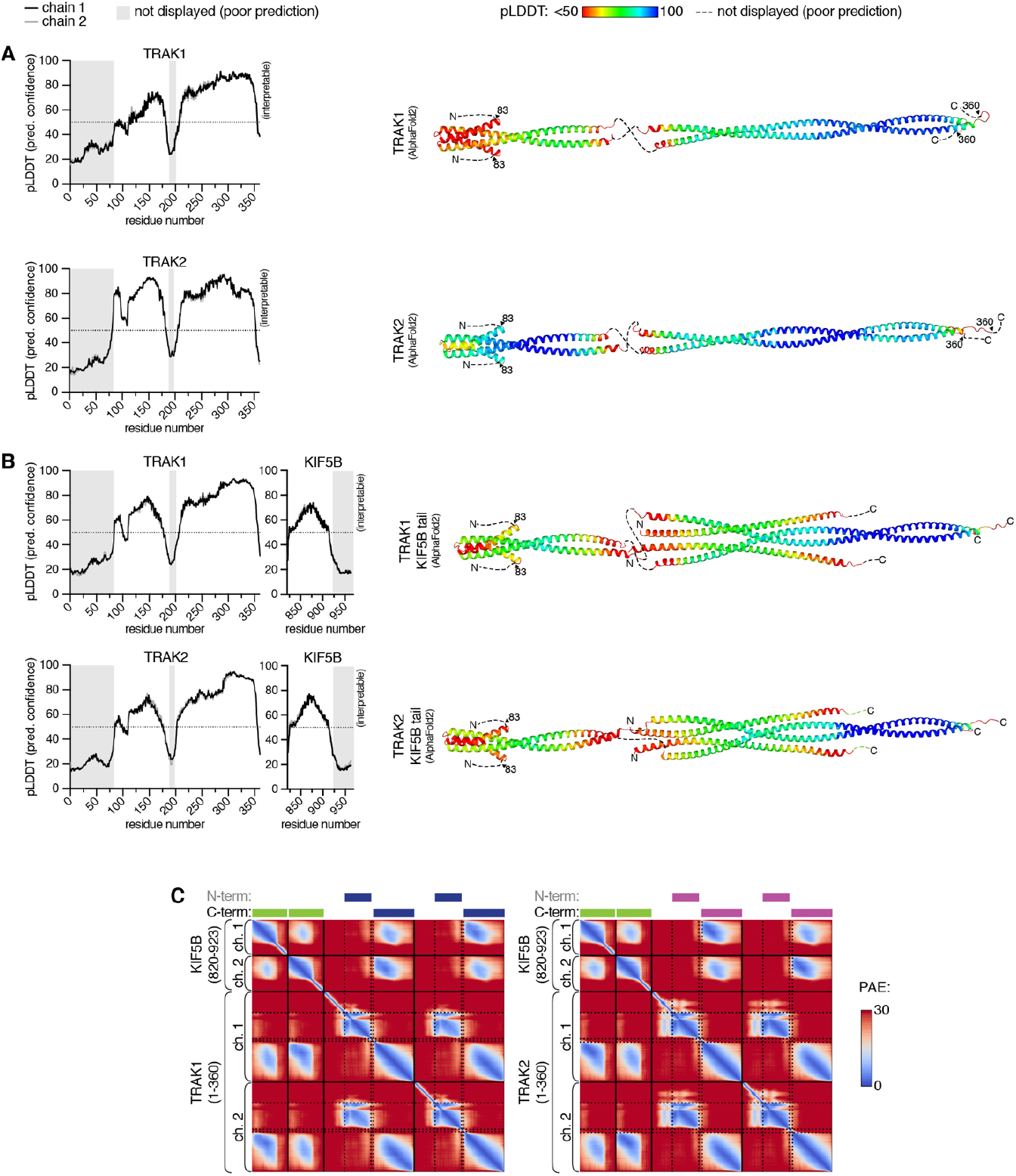
AlphaFold2 prediction confidence. (**A**) AlphaFold2-derived pLDDT scores for the top-ranked predictions of the N-terminal coiled-coil dimers of TRAK1 (top) and TRAK2 (bottom). Left: per-residue pLDDT plots for each coiled-coil prediction. Modelled residues with low confidence scores are highlighted using a grey bar, and are not displayed. Right: per-residue pLDDT prediction confidence mapped onto the predicted structures. For clarity, the coiled-coil is split into two separately aligned sections at the site of disorder around aa 200. (**B**) AlphaFold2-derived pLDDT scores for the top-ranked predictions of the N-terminal coiled-coil dimers of TRAK1 (top) and TRAK2 (bottom) in complex with kinesin-1. Left: per-residue pLDDT plots for each complex prediction. Modelled disordered residues with low confidence scores are highlighted using a grey bar, and are not displayed. Right: per-residue pLDDT prediction confidence mapped onto the predicted structures. For clarity, the coiled-coil is split into two separately aligned sections at the site of disorder around aa 200. (**C**) AlphaFold2-derived PAE plots for the top-ranked predictions of the N-terminal coiled-coil dimers of TRAK1 (top) and TRAK2 (bottom) in complex with kinesin-1. The N- and C-terminal portions of the structures displayed in (B), Fig. 2A (top), and fig. S4A (top) are indicated across the top of the plot.

**Fig. S6.**
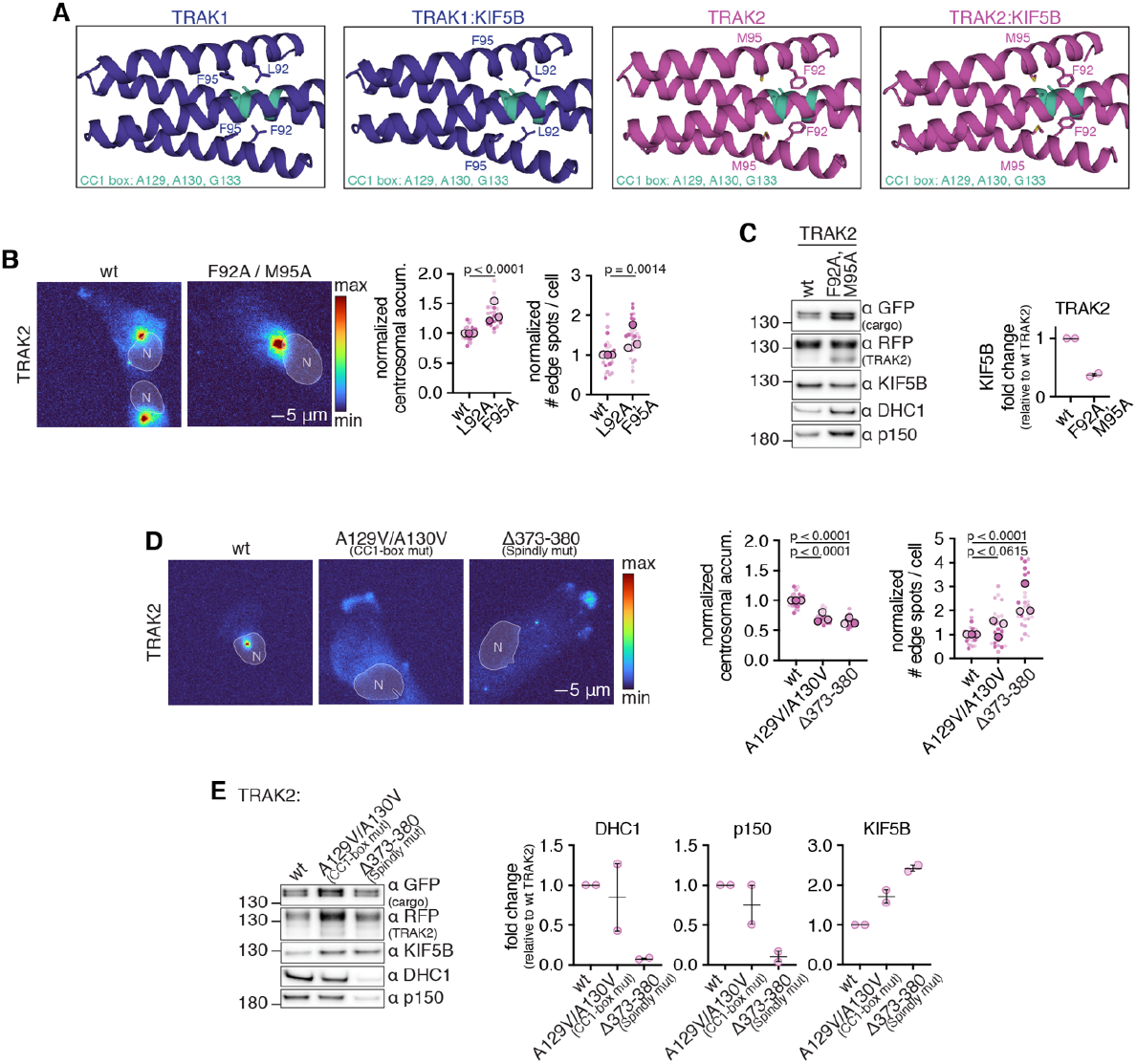
Probing the AlphaFold2 model for dynein repression. (**A**) Insets from the AlphaFold2 predictions shown in Fig. 2A and fig. S4A showing the N-terminal helix folded back onto the CC1-box region (cyan). (**B**) Left: Representative maximum intensity projection images of cargo from cell lines expressing wt or L92A/F95A TRAK2. Right: Phenotype quantification from three biological replicates as shown on the left. For perinuclear accumulation n(wt)=27; n(L92A/F95A)=27; p<0.0001 (two-tailed t-test, t=7.898, df=52). For peripheral accumulation n(wt)=26; n(L92A/F95A)=26; p=0.0014 (two-tailed t-test, t=3.390, df=50). (**C**) Left: Immunoblot analysis of the motor-adaptor components associated with FLAG-purified cargo from indicated cell lines. Right: quantification of relative motor protein levels normalized to wt TRAK2. Error bars show the median and 95% confidence interval; n = 2 for all conditions. (**D**) Left: Representative maximum intensity projection images of cargo from cell lines expressing wt, A129V/A130V (CC1-box mutant) or Δ373-380 (Spindly motif mutant) TRAK2. Right: Phenotype quantification from three biological replicates as shown on the left. For perinuclear accumulation n(wt)=27; n(A129V/A130V)=27; p<0.0001 (two-tailed t-test, t=10.46, df=52); n(Δ373-380)=26; p<0.0001 (two-tailed t-test, t=13.35, df=51). For peripheral accumulation n(wt)=26; n(A129V/A130V)=25; p<0.0615 (two-tailed t-test, t=1.914, df=49); n(Δ373-380)=26; p<0.0001 (two-tailed t-test, t=6.358, df=50). (**E**) Left: Immunoblot analysis of the motor-adaptor components associated with FLAG-purified cargo from indicated cell lines. Right: quantification of relative motor protein levels normalized to wt TRAK2. Error bars show the median and 95% confidence interval; n = 2 for all conditions.

**Fig. S7.**
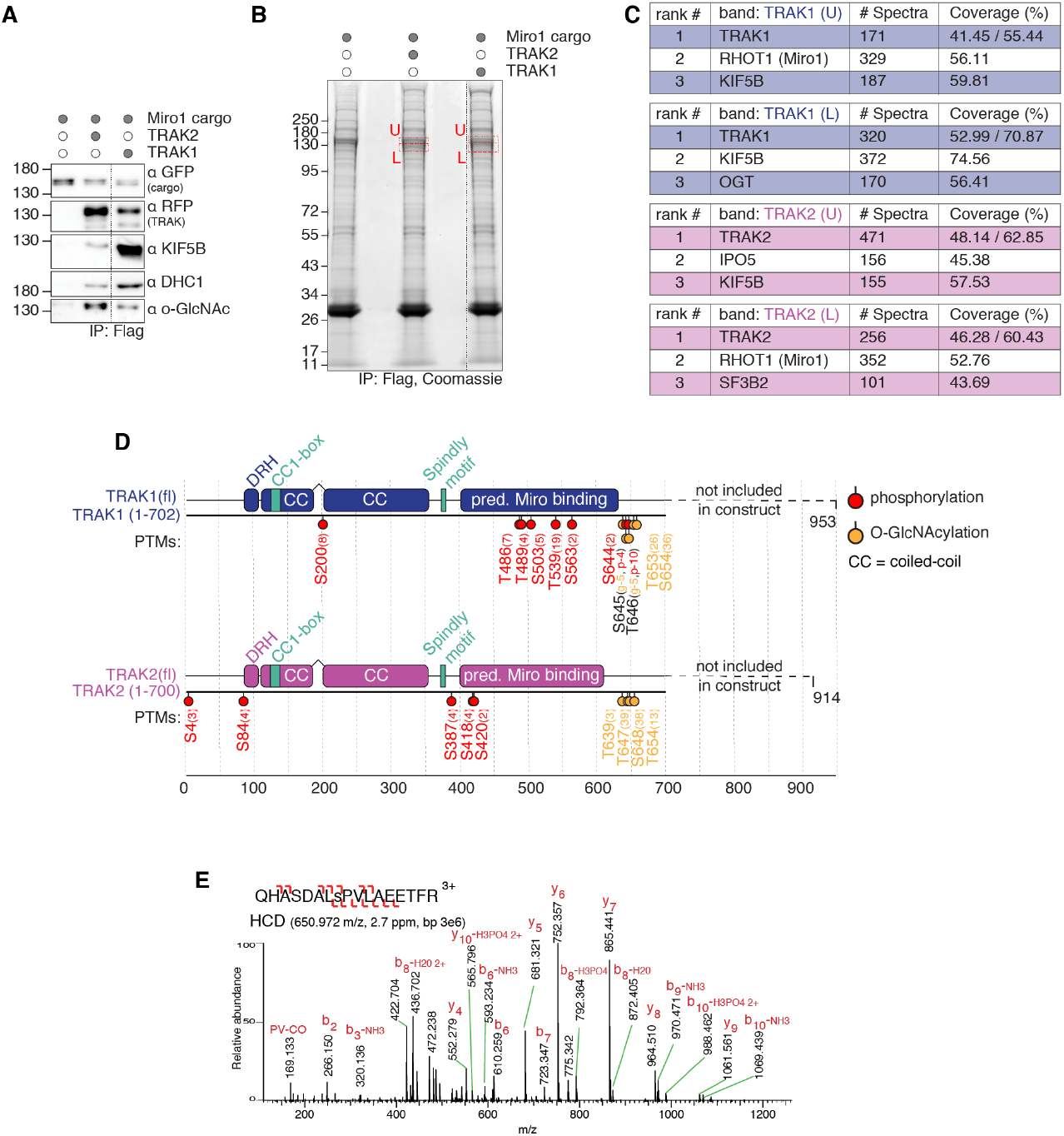
Mapping post-translational modifications of TRAK. (**A**) Immunoblot analysis of the FLAG-purified cargo used for mass spectrometry. (**B**) SDS-PAGE separation of co-eluting proteins from purification analysed in (A). Bands cut for further protein and PTM analysis are highlighted in red. (**C**) Top ranked proteins identified in each sample. For TRAK, coverage is reported as % of full-length sequence / overexpressed construct. (**D**) PTM sites identified from bands cut in (B). Peptide spectra matches for each site are reported in brackets. (**E**) HCD mass spectrum for the TRAK2 phosphopeptide demonstrating modification of S84, in lower case. Observed m/z, mass error in ppm, and base peak (bp) is shown. The TRAK2 S84 phosphorylation site was additionally found in several high-throughput studies (in PhosphoSitePlus(*49–51*), while no phosphorylation of T84 on TRAK1 has been reported (in PhosphoSitePlus).

**Fig. S8.**
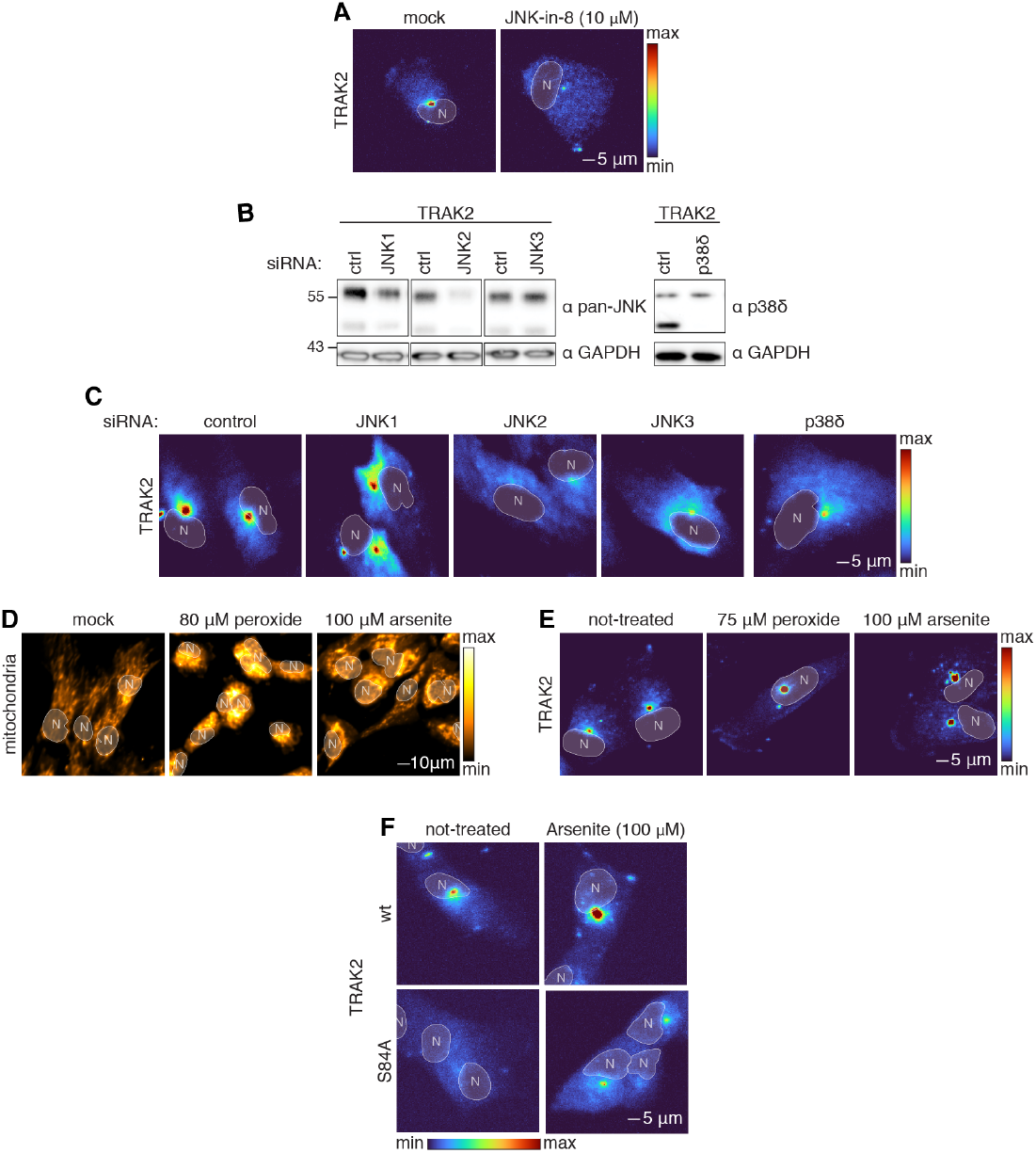
Characterizing Ser84 phosphorylation of TRAK2. Quantification corresponding to panels A, C, D, E, F is shown in Fig. 3 E, F, H, I, J respectively. (**A**) Representative maximum intensity projection images of cargo from cell lines expressing TRAK2 following mock or JNK-in-8 (10 μM) treatment (3h). (**B**) Immunoblot analysis of whole cell lysate showing the efficacy of siRNA treatments used in Fig. 3F and (C). (**C**) Representative maximum intensity projection images of cargo from cell lines expressing TRAK2 following RNAi treatment as indicated. (**D**) Representative maximum intensity projection images of mitochondria in untransduced cells following oxidative treatment as indicated. (**E**) Representative maximum intensity projection images of cargo from cell lines expressing TRAK2 following oxidative treatment as indicated. (**F**) Representative maximum intensity projection images of cargo from cell lines expressing wt or S84A TRAK2 following arsenite treatment.

**Movie S1. Synthetic cargo accumulation is microtubule dependant**.

Reduction in accumulation of (**A**) TRAK1- and (**B**) TRAK2-driven synthetic cargo following microtubule disassembly by nocodazole treatment (2.5 μM, 30 min), and re-accumulation following microtubule re-assembly mediated by nocodazole wash-out. Scale bar = 5 μm. Time = mm:ss.

**Movie S2. Synthetic cargo dynamics**.

Dynamics of PA JF-646-labelled cargo after photoactivation of synthetic cargo in the absence of TRAK co-expression (**A**), TRAK1 co-expression (**B**), TRAK2 co-expression (**C**). The video for each condition is preceded by a still image showing cell orientation. Photoactivatable JF-646-labelled cargo dynamics mediated by TRAK expression as indicated. Scale bar = 5 μm. Time = mm:ss.

## Notes

### Competing Interest Statement

The authors have declared no competing interest.

